# DDO-adjuvanted influenza A virus nucleoprotein mRNA vaccine induces robust humoral and cellular type 1 immune responses and protects mice from challenge

**DOI:** 10.1101/2024.10.27.620508

**Authors:** Victoria Gnazzo, Hanaa Saleh, Ítalo Castro, Adrianus C.M. Boon, Amelia K Pinto, James D. Brien, Carolina B. López

## Abstract

A challenge in viral vaccine development is to produce vaccines that generate both neutralizing antibodies to prevent infection and cytotoxic CD8^+^ T-cells that target conserved viral proteins and can eliminate infected cells to control virus spread. mRNA vaccines offer an opportunity to design vaccines based on conserved CD8-targeting epitopes, but achieving robust antigen-specific CD8^+^ T-cells remains a challenge. Here we tested the viral-derived oligonucleotide DDO268 as an adjuvant in the context of a model influenza A virus (IAV) nucleoprotein (NP) mRNA vaccine in C57BL/6 mice. DDO268 safely induced local type I interferon (IFN) production, stimulated dendritic cells type 1 (DC1) activation and migration to the draining lymph nodes, and improved the generation of IgG2c antibodies and antigen-specific effector Th1 CD4^+^ and CD8^+^ T-cells (IFNγ^+^TNFα^+^IL2^+^) when co-packaged with NP mRNA. The DDO268 adjuvanted vaccine provided enhanced protection against lethal IAV challenge and reduced the antigen dose required to achieve this protection. These results highlight the potential of DDO268 as an effective mRNA vaccine adjuvant and show that an IAV NP mRNA/DDO268 vaccine is a promising approach for generating protective immunity against conserved IAV epitopes.

**IMPORTANCE:** Vaccines that generate neutralizing antibodies and cytotoxic CD8^+^ T-cells targeting conserved epitopes are ideal for effective protection against viruses. mRNA vaccines combined with the right adjuvant offer a promising solution to this challenge. We show that the virus-derived oligonucleotide DDO268 enhances antibody and T cell responses to an influenza A virus (IAV) nucleoprotein (NP) mRNA vaccine in mice. DDO268 safely induces local type I interferon production and stimulates dendritic cell activation providing enhanced protection against IAV challenge. In addition, the adjuvant activity of DDO268 allows for the use of lower antigen doses during vaccination.

## INTRODUCTION

The ideal antiviral vaccine would induce immunity against the primary target virus and its variants offering “broad” or “universal” protection. This vaccine would elicit virus-neutralizing antibodies to prevent infection and would generate type 1 cellular immunity, including antigen-specific Th1 CD4^+^ T-cells and CD8^+^ T-cells that recognize conserved viral antigens, eliminate infected cells, and prevent reinfection.

mRNA vaccines represent a milestone in vaccinology, offering safety, flexibility, cost-effective manufacturing, and rapid development [1, 2]. mRNA vaccines facilitate antigen selection, allowing for easier development of viral vaccines that target conserved CD8^+^ T-cells epitopes. However, in addition to the right antigen, proper immune system direction is needed for stimulating a type 1 immune response. Type I interferons (IFNs) are key drivers of antigen-specific Th1 CD4^+^ T-cells and cytotoxic CD8^+^ T-cells during infection [3–6]. In the context of vaccines, immunity can be directed by including type I IFN-inducing molecules.

mRNA vaccines consist of the mRNA encoding the target protein and the lipid nanoparticle (LNP) that protects the mRNA from degradation. The LNP component induces expression of chemokines and cytokines that assist with activating and recruiting immune cells to the inoculation site and initiate the adaptive immune response [7]. The mRNA component can also provide immune-activating signals, particularly when byproducts of the mRNA *in vitro* transcription (IVT) are present [8]. These byproducts can form double-stranded (ds)RNA structures that bind to cellular RNA sensor proteins, triggering the expression of type I IFNs. However, to minimize excessive inflammation and adverse reactions caused by dsRNA byproducts [9], human mRNA vaccines are either prepared using modified nucleotides to limit their binding to RNA sensor molecules, or the mRNA is purified eliminating dsRNA products that trigger the expression of type I IFNs [10].

Considering the above, the immunostimulatory properties of mRNA vaccines can be improved by including specific type I IFN-inducing molecules that can be tittered for controlled type I IFN induction. Our previous work identified non-standard viral genome-derived oligonucleotides (DDOs) as effective triggers of antigen-specific type 1 immune responses during vaccination with inactivated or purified viral proteins [11, 12]. DDOs are synthetic RNAs derived from the 546 nt-long Sendai virus (SeV) copy-back viral genome and activate cellular RNA viral sensors leading to type I IFN production [12]. Here we explore the use of DDOs as a type-1 immunity inducing adjuvant for an influenza A virus (IAV) model vaccine.

IAV causes approximately 3–5 million severe disease cases and 290,000–650,000 deaths annually [13]. Yearly influenza epidemics and periodic pandemics result from the virus constant antigenic variation, which helps it evade host immune responses. Current IAV vaccines offer high levels of strain-specific protection but are less effective against new, antigenically distinct strains requiring frequent reformulation [14]. To address this problem, in addition to targeting neutralizing antigens exposed in the cell surface, vaccines could be directed to the internal viral nucleoprotein (NP) that is largely conserved among IAVs [15–18] and contains epitopes that are primary antigenic targets of T cells (8).

As a proof of concept for using DDO as a type-I IFN-inducing adjuvant in mRNA vaccines targeting conserved viral antigens, we developed a DDO268-adjuvanted mRNA vaccine encoding the IAV NP. This vaccine induced localized type I IFN production leading to the activation of NP-specific CD4^+^ and CD8^+^ T-cells and providing enhanced protection against IAV than the NP mRNA vaccine alone. Overall, DDO268 significantly enhanced the immune response to IAV NP in the context of an mRNA vaccine by providing controlled induction of type I IFN necessary for robust T cell responses.

## RESULTS

### *In vivo* administration of DDO268 is safe and elicits a localized immune response without systemic effects

Our previous work [12] demonstrated that DDO268 effectively induces expression of type I interferon (IFN) at the site of inoculation. To further assess the safety of DDO268 as an adjuvant, we inoculated C57BL/6 mice subcutaneously in the footpad with 50 µg of DDO268, a dose 10 times higher than that used in our previous studies [11, 12]. We evaluated several parameters, including complete blood counts (CBC), serum chemistry, and systemic cytokine levels, up to 72 hours post-inoculation. As shown in **Fig. 1** and **Table S1**, DDO268 did not induce significant changes in any of the measured parameters. Analysis of the leukogram, erythrogram, and thrombocyte counts (**Fig. 1A-G**) revealed no significant alterations in blood cell populations. Similarly, chemical parameters such as blood urea nitrogen, bilirubin, and aspartate transferase levels (**Fig. 1H-J**) were comparable to those in mice inoculated with PBS. Additionally, transcripts of pro-inflammatory cytokines, including *Il6*, *Mx1*, and *TNF*α, were not detected in the liver at any time point (**Fig. 1K-M**). These findings support the safety profile of DDO268, confirming its ability to elicit a localized immune response without systemic effects.

**Figure 1:**
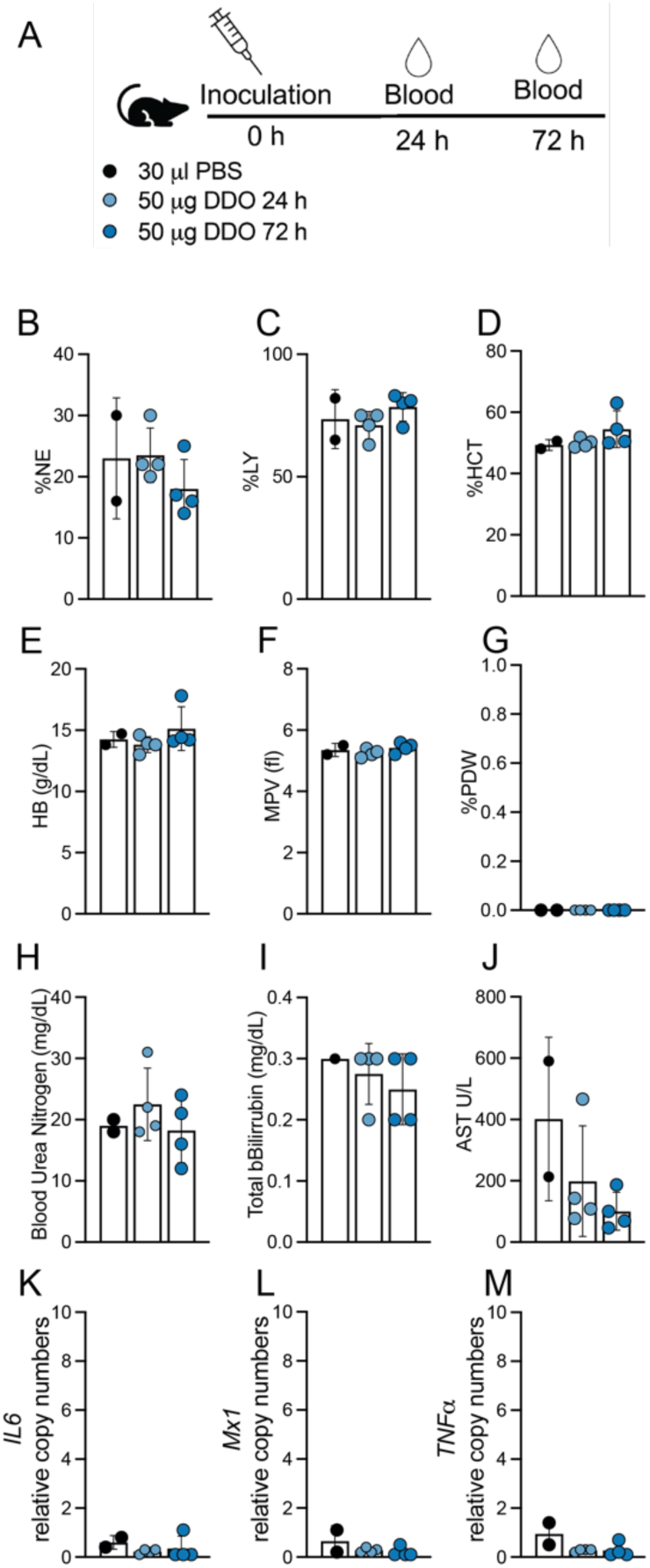
Absence of systemic toxicity after DDO268 inoculation in the footpad. (**A**) Study design. C57BL6 mice were inoculated s.c. with 50µg DDO in the rear footpad. (**B-C**) Leukogram results for percent neutrophils (%NE) and leukocytes (%LY). (**D-E**) Erythrogram results for percent hematocrit (%HCT) and hemoglobin concentration (g/dL) (HB g/dL). (**F-G**) Thrombocytes showing mean platelet volume (femtoliters)(MPV fl) and Platelet Distribution Width (% PDW). (**H-J**) Chemical parameters measurements: Blood Urea Nitrogen (BUN mg/dL); Total Bilirubin (mg/dL); Aspartate Aminotransferase (AST U/L). (**K-M**) Transcript levels of *IL6, Mx1* and *TNFα* relative to the housekeeping genes GAPDH and *β-actin* in liver of inoculated mice. The mean ± SD of each group is shown n = 2 (mock) and 4(treated).

### DDO268 promotes the generation of IgG2c antibodies and antigen-specific CD8^+^ T-cells in response to a SARS-CoV-2 mRNA vaccine

To evaluate DDO268 in the context of the mRNA vaccine platform, we tested its impact on the induction of type 1 immune responses during vaccination with the original SARS-CoV-2 mRNA vaccine, BNT162b2. We vaccinated C57BL6 mice subcutaneously in the footpad with a mix of 0.125 µg BNT162b2 vaccine and 5 µg DDO268, a vaccine dose that showed no local inflammation, and boosted the mice with the same dose 28 days later (**Fig. 2A**). As we showed that migration of conventional dendritic cells type 1 (cDC1) to the draining lymph node is an essential early step in the generation of type 1 immune responses upon immunization in the presence of DDO268 [12], we first assessed cDC1 migration to the popliteal lymph node upon vaccination with either DDO268 alone, BNT162b2 alone, BNT162b2 combined with an inert RNA (a synthetic 30 nucleotide-long and non-immunostimulatory RNA), or BNT162b2 combined with DDO268. All combinations were used at equivalent molar amounts. DDO268, but not the non-immunostimulatory RNA, significantly increased the number of cDC1 in the draining lymph nodes at 12 hours post vaccination (**Fig. 2B** and **Fig. S1)**, a previously determined peak timepoint for cDC1 accumulation in the lymph node upon vaccination [12].

**Figure 2:**
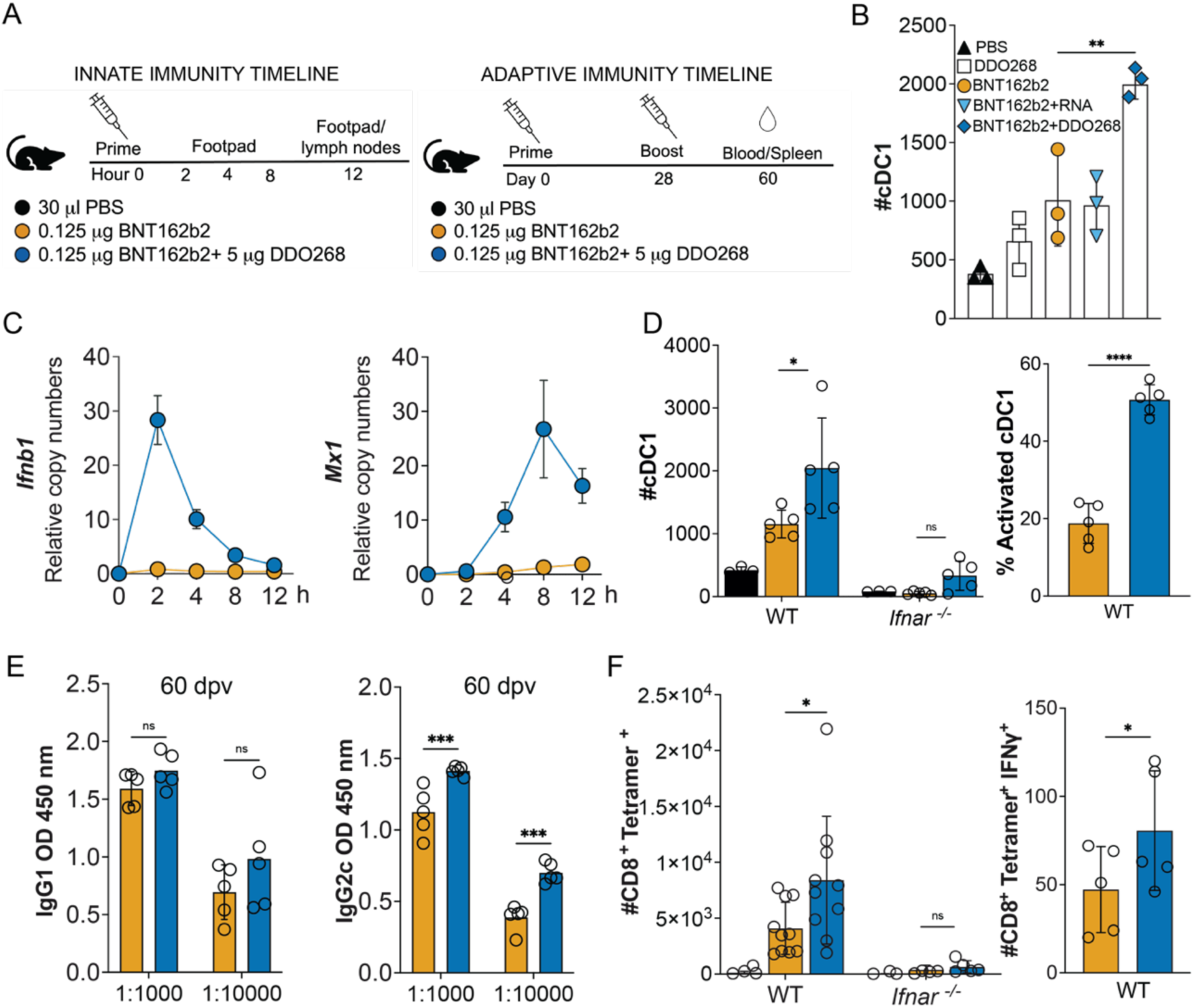
DDO268 improves type 1 immune responses to the original Pfizer SARS-CoV2 mRNA vaccine. C57BL6 WT and *Ifnar^-/-^* mice were immunized in the rear footpad with the Pfizer SARS-CoV-2 mRNA vaccine BNT162b2 (0.125 µg), or with BNT162b2 (0.125 µg) + DDO268 (5 µg). (**A**) Timeline and groups for the study design. (**B**) Number of total cDC1 in the draining lymph nodes 12 hours after vaccination of C57BL6 WT mice with: PBS, DDO268 (5 µg), BNT162b2 (0.125 µg), BNT162b2 (0.125 µg) + non-immunostimulatory RNA (5 µg) or BNT162b2 (0.125 µg) + DDO268 (5 µg). cDC1 were characterized as live, CD3^-^NK1.1^-^B220^-^CD19^-^MHCII^hi^CD64^-^Ly6c^-^CD11c^hi^XCR1^+^SIRPa^-^. N = 3 mice per group (**C**) *Ifnb1* and *Mx1* transcript level at the inoculation site of WT mice measured by qPCR at 2, 4, 8, and 12 hours after vaccination. (**D**) Number of total and activated cDC1 in the draining lymph nodes 12 hours after vaccination of WT and *Ifnar^-/-^* mice. cDC1 were characterized as Live, CD3^-^NK1.1^-^B220^-^ CD19^-^ MHCII^hi^CD11c^hi^ CD64^-^Ly6c^-^ XCR1^+^ SIRPa^-^, activated cDC1 are Live, CD3^-^NK1.1^-^B220^-^CD19^-^MHCII^hi^CD11c^hi^ CD64^-^Ly6c^-^ XCR1^+^ SIRPa^-^ CD86^+^ (**E**) SARS-CoV-2 Spike specific IgG21 and IgG2c antibodies in vaccinated animals 60 days post vaccination (32 days post booster). (**F**) Number of CD8^+^ Tetramer^+^ T-cells in the spleens on day 39 after boost and specific CD8^+^ Tetramer^+^ IFNγ^+^ T-cells in the spleen. Number of cells shown was normalized to 500,000 live cells. In all experiments the mean ± SEM of each group is shown (n = 3-5/group) except for (F) WT # CD8^+^ Tetramer^+^ where data was pooled from two independent experiments n= 5 mice each. *=p <0.05, **=p <0.01, ***=p <0.005 ****=p <0.001 by unpaired t-test for comparisons between two groups, and one-way ANOVA or two-way ANOVA with Bonferroni’s multiple comparison test for comparisons among three or more groups.

To confirm the induction of type I IFNs at the site of immunization, we measured type I IFN gene *Ifnb1* and the IFN-stimulated gene *Mx1* in the footpad after vaccine administration. As shown in **Fig. 2C**, DDO268 transiently and significantly enhanced the expression of these genes. In addition, in mice lacking the type I IFN receptor (*Ifnar* KO), cDC1 migration was abolished, underscoring type I IFN’s role in the response to DDO268 (**Fig. 2D**). Moreover, DDO268 increased titers of circulating IgG2c antibodies (**Fig. 2E**) and promoted the generation of a larger population of activated SARS-CoV-2 specific CD8^+^ IFNγ^+^ T-cells in the spleen compared to BNT162b2 alone (**Fig. 2F**). The adaptive immune response enhancement by DDO268 depended on type I IFN signaling, similar to our previous findings with inactivated virus vaccines [12]. Overall, these proof of principle experiments suggest that DDO268 can enhance the immune response elicited by an mRNA vaccine toward the generation of type 1 humoral and cellular immunity in a type I IFN-dependent manner.

### Development and evaluation of an IAV NP mRNA/DDO268 vaccine

We next investigated if DDO268 promote type 1 immunity when co-packaged with the mRNA. To this end, we developed a model mRNA vaccine using the IAV nucleoprotein (NP) as target antigen. To eliminate unspecific immunostimulatory dsRNA, we cellulose-purified the *in vitro* transcribed unmodified IAV NP mRNA followed by stringent quality control (**Fig. 3A-B**). Codon-optimized influenza A/Puerto Rico/8/1934 H1N1 NP mRNA was generated from the pJB201.1 vector (**Fig. 3A**) and the mRNA functionality tested in A549 cells. No differences in NP expression were observed between purified and unpurified mRNAs within the first 48 hours post-transfection (**Fig. 3C**), but significant differences in *Ifnb1* expression were observed with the purified mRNA inducing no significant cytokine expression (**Fig. 3D**). In all subsequent experiments we used unmodified dNTPs for mRNA production followed by cellulose-purification.

**Figure 3:**
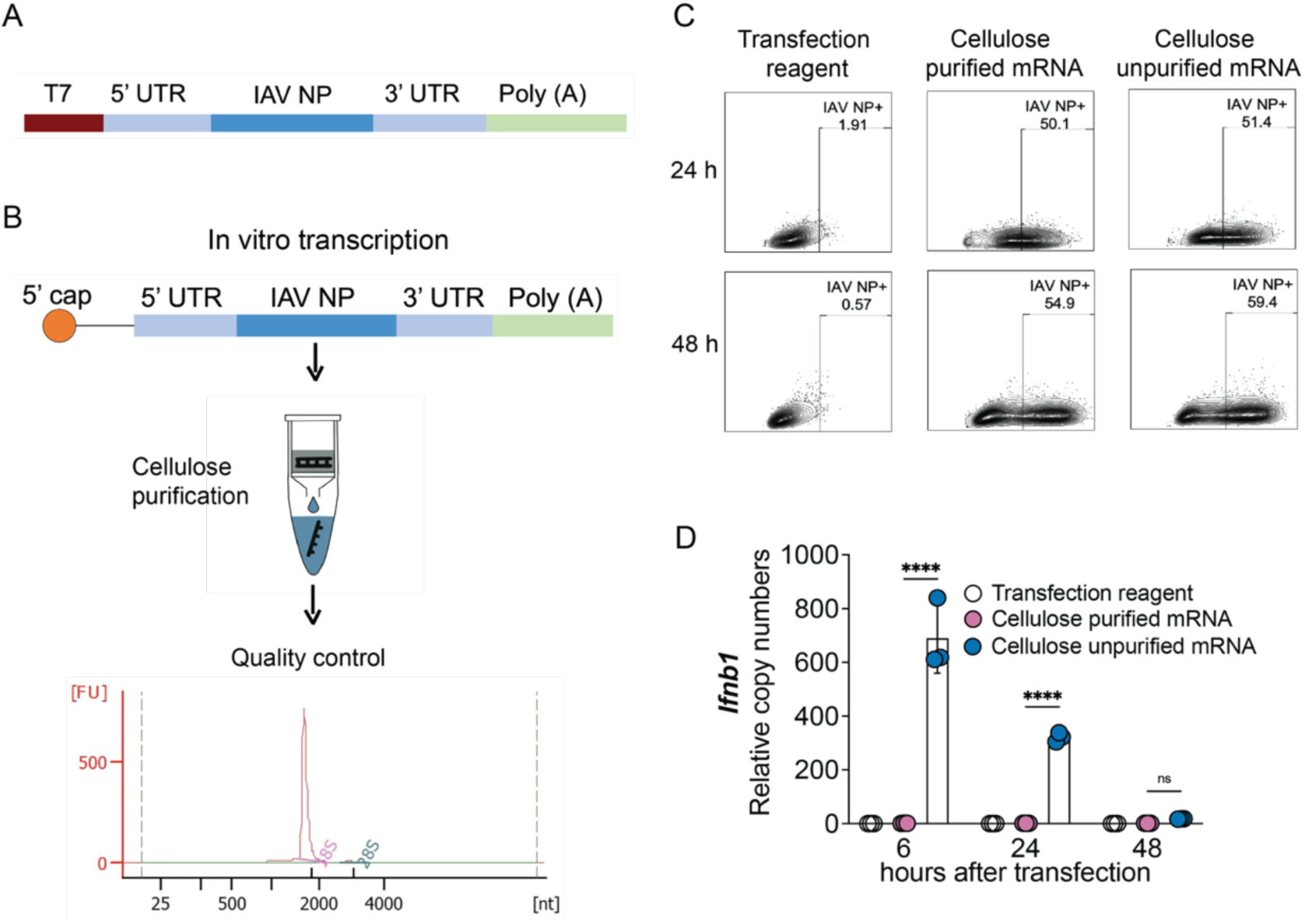
Design, characterization and functional assessment of IAV NP mRNA. (**A**) pJB201.1 plasmid schematic. (**B**) mRNA encoding the IAV NP *in vitro* transcription and downstream process. Cellulose purification and quality control by Bioanalyzer. (**C, D**) Functional assessment of cellulose purified and unpurified mRNA. **(C)** Comparison of IAV NP expression in A549 cells (1x10^6^ cells) transfected with 1mg of cellulose purified or unpurified IVT NP mRNA at 24 and 48 hours post transfection. Protein detection by flow cytometry upon intracellular staining with anti-NP antibody. **(D)** Transcript levels of *Ifnb1* relative to the housekeeping genes GAPDH and *β-actin* in A549 cells at 6, 24 and 48 hours after transfection with cellulose purified and unpurified IVT NP mRNA. The mean ± SD of each group is shown (n = 3/group) ****=p <0.001 by one-way ANOVA test.

We formulated mRNA vaccines into LNPs using Gen-Voy ILM^TM^ reagent and co-packaged the IAV NP mRNA with a series of immunostimulatory and non-immunostimulatory RNAs. The formulations included: IAV NP mRNA alone, IAV NP mRNA/DDO268, IAV NP mRNA/DDO268B (a modified DDO268 molecule, with the immunostimulatory motif encoded in reverse direction than DDO268), IAV NP mRNA/X region (a non-immunostimulatory small RNA derived from the hepatitis C virus genome [19, 20]), and empty LNPs (**Fig. 4A**). Each formulation contained a 1:1 molar ratio of mRNA to RNA, ensuring consistent total number of RNA molecules. The RNA concentration and the encapsulation efficiency were assessed using RiboGreen RNA assay [21], and formulation sizes (ranged from 120 to 150 nm) were measured by Dynamic Light Scattering (DLS) (**Fig. 4B**).

**Figure 4:**
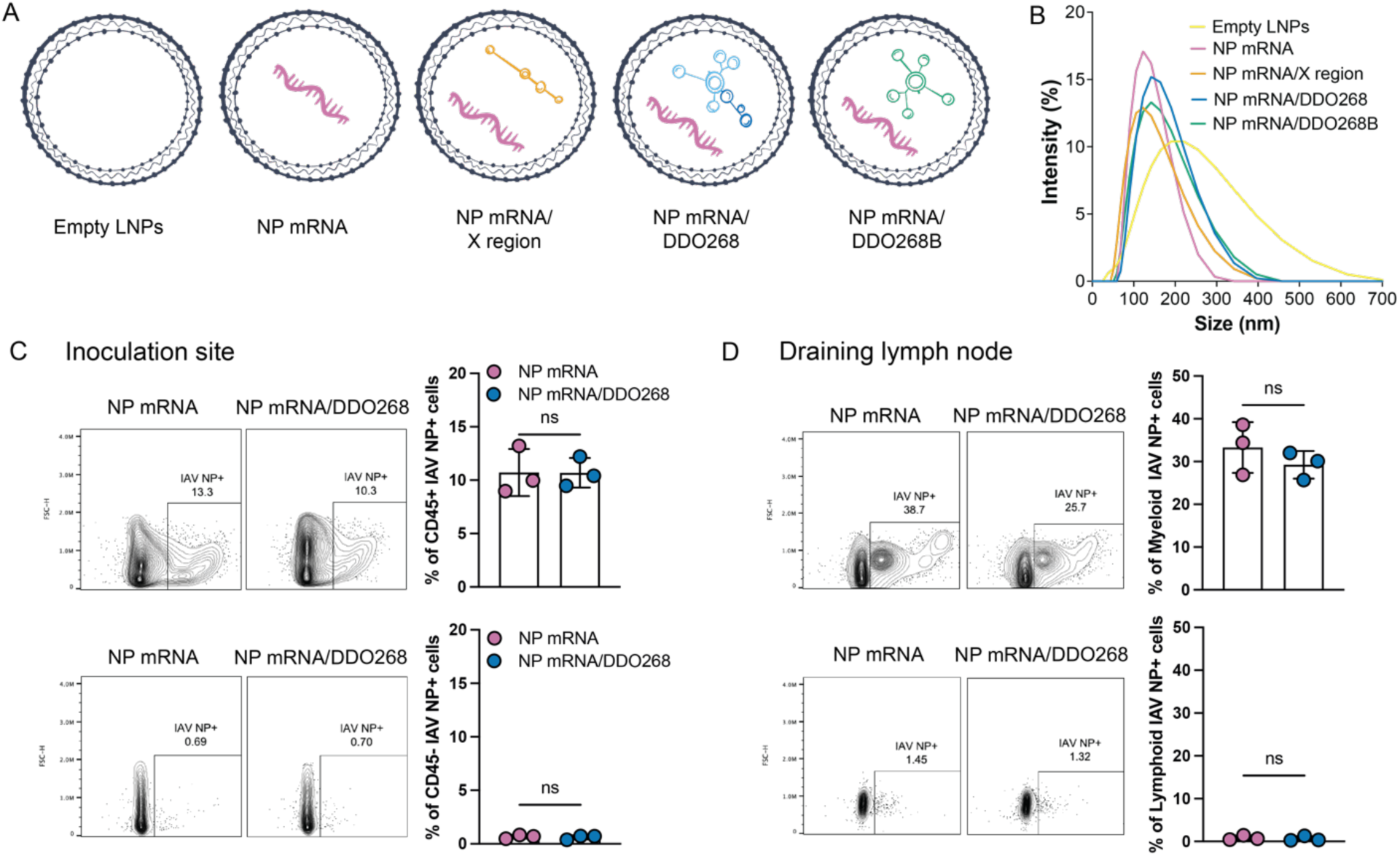
IAV NP mRNA vaccine formulation, characterization and *in vivo* testing. **(A)** Scheme of the different vaccine formulations tested: Empty LNP; IAV NP mRNA; IAV NP mRNA/X region; IAV NP mRNA/DDO268; IAV NP mRNA/DDO268B (**B**) Size distribution of nanoparticles empty or carrying IAV NP mRNA; IAV NP mRNA/X region; IAV NP mRNA/DDO268; IAV NP mRNA/DDO268B measured by Dynamic Light Scattering. (**C, D**) Expression of IAV NP at the **(C)** inoculation site (rear footpad) or **(D)** draining lymph nodes detected by flow cytometry upon intracellular staining with anti-NP antibody 24 hours after vaccination of C57B6 mice with 1 µg NP mRNA alone or 1 µg or NP mRNA/DDO268. Gating for immune cells (CD45^+^): single cells, live, CD45^+^ IAV NP^+^. Gating for non-immune cells (CD45^-^): single cells, live, CD45^-^ IAV NP^+.^ Gating for myeloid cells: single cells, live, CD3^-^ B220^-^ CD19^-^ NK1.1^-^ IAV NP^+^. Gating for lymphoid cells: single cells, live, CD3^+^ B220^+^ CD19^+^ NK1.1^+^ IAV NP^+^. Gating for myeloid cells: single cells, live, CD3^-^ B220^-^ CD19^-^ NK1.1^-^ IAV NP^+^. The mean ± SD of each group is shown n = 3/group ns=p > 0.05 by unpaired t-test.

To assess protein expression and identify the cell types expressing the viral protein *in vivo*, we inoculated mice subcutaneously and evaluated NP expression at the inoculation site and in the draining lymph nodes at 24 hours post-vaccination. NP was expressed in CD45^+^ cells in the footpad (**Fig. 4C**) and in myeloid cells within draining lymph nodes (**Fig. 4D**). These results agree with previous reports of mRNA vaccines being primarily taken up by immune cells at the site of immunization [22–24].

### DDO268 confers type I IFN-inducing ability to the IAV NP mRNA vaccine

To confirm that our vaccine induced a local expression of type I IFN, cytokines and chemokines, we analyzed footpads and popliteal lymph nodes at 12 hours after subcutaneous inoculation, as well as spleens at 12 and 24 hours after subcutaneous inoculation (**Fig. 5A**). As expected, empty LNPs induced *Il1b*, *Il6* and *Ccl2* expression [7] but showed minimal *Ifnb1* or *Cxcl10* expression at the inoculation site (**Fig. 5B**). IAV NP mRNA and IAV NP mRNA/X region formulations induced low *Ifnb1* levels, while the IAV NP mRNA/DDO268B formulation showed higher levels than controls but lower than the DDO268 formulation (**Fig. 5C**). All vaccine formulations induced *Il6*, *Il1b, Ccl2*, and *Cxcl10* transcription (**Fig. 5D-G**). *Il13* transcripts a type 2 immunity-associated cytokine, were undetected at the inoculation site with any formulation (**Fig. 5H**). Importantly, DDO268-containing formulations did not induce significant *Ifnb1* in the spleen at 12 hours after vaccination (**Fig. 5I**) or *Il6*, Mx1, and TNF expression in the spleen at 24 hours after vaccination (**Fig. 5J-L**) confirming that DDO268 adjuvancy is safe as it induced a localized immune response [12]. Additionally, cDC1 recruitment to the draining lymph nodes increased significantly with DDO268 formulations (**Fig. 5M**).

**Figure 5:**
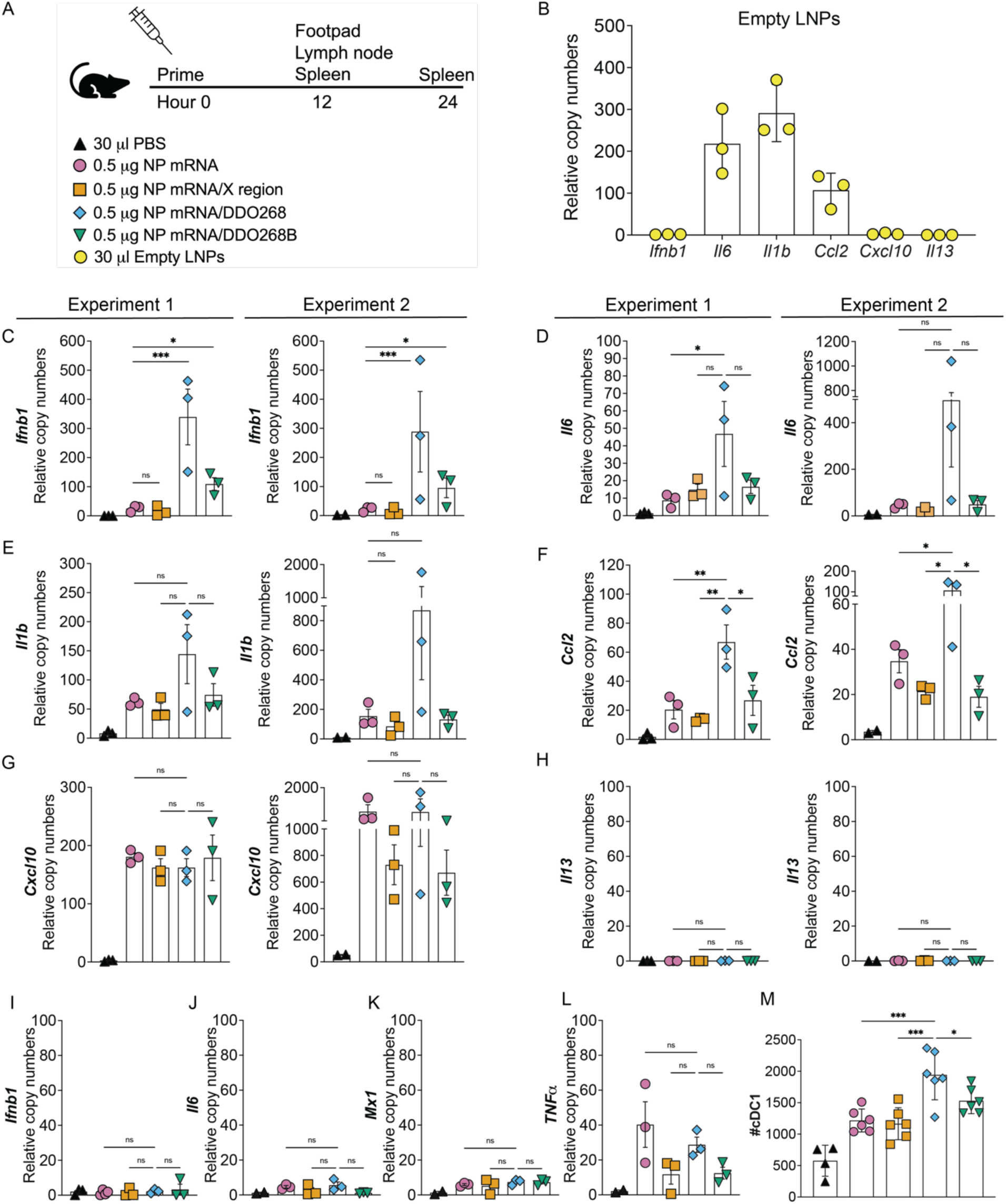
DDO268 promotes strong type 1 innate immune responses to the IAV NP mRNA vaccine. **(A)** Timeline and groups for the study design. **(B)** Transcript levels of *Ifnb1*, *Il6, Il1b*, *Ccl2*, *Cxcl10* and *Il13* relative to the housekeeping genes GAPDH and *β-actin* in the footpad of C57BL6 mice 12 hours after vaccination with empty LNPs. Transcript levels of **(C)** *Ifnb1*, **(D)** *Il6,* **(E)** *Il1b*, **(F)** *Ccl2*, **(G)** *Cxcl10* and **(H)** *Il13* relative to the housekeeping genes GAPDH and *β-actin* in the footpad of C57BL6 mice 12 hours after vaccination with PBS; 0.5 µg NP mRNA or 0.5 µg NP mRNA/X region; 0.5 µg NP mRNA/DDO268; 0.5 µg NP mRNA/DDO268B (**I)** Transcript levels of *Ifnb1* in spleen of C57BL6 mice 12 hours after vaccination with PBS; 0.5 µg NP mRNA or 0.5 µg NP mRNA/X region; 0.5 µg NP mRNA/DDO268; 0.5 µg NP mRNA/DDO268B. Transcript levels of **(J)** *Il6,* **(K)** *Mx1,* and **(L)** *TNF*α in spleen of C57BL6 mice 24 hours after vaccination with PBS; 0.5 µg NP mRNA or 0.5 µg NP mRNA/X region; 0.5 µg NP mRNA/DDO268; 0.5 µg NP mRNA/DDO268B. **(M)** Number of cDC1 in the draining lymph nodes 12 hours after vaccination with PBS; empty LNPs; 0.5 µg NP mRNA or 0.5 µg NP mRNA/X region; 0.5 µg NP mRNA/DDO268; 0.5 µg NP mRNA/DDO268B. cDC1 were characterized as Live, CD3^-^NK1.1^-^B220^-^CD19^-^ MHCII^hi^CD11c^hi^ CD64^-^Ly6C^-^ XCR1^+^SIRPa^-^. Mean ± SEM of each group is shown (n = 3/group) *=p <0.05; **=p <0.01; ***=p <0.005, by one-way ANOVA with Bonferroni’s multiple comparison test. (**C-H**) Two independent experiments are shown n=3 mice each. (**M**) Data represent two independent experiments where data was pooled with n=3 mice, n = 2/3 for PBS group.

These findings highlight three key insights: (1) cells recognize and respond to DDO268 within lipid nanoparticle, (2) DDO268 induces type I IFN expression necessary to initiate a type 1 immune response by mobilizing cDC1s to the draining lymph node [12] and (3) DDO268 induces a localized inflammatory response.

### IAV NP mRNA/DDO268 vaccine enhances antigen-specific type 1 humoral and cellular immune responses

We next evaluated the adaptive immune responses elicited by the IAV NP mRNA/DDO268 vaccine after primary vaccination and a booster administered four weeks later (**Fig. 6A**). We also tested DDO268B, which triggers less robust responses **(Fig. 5** and **Fig. S2**). Three weeks post-booster, anti-NP antibody levels were measured from sera, including specific IgG1 and IgG2c isotypes to determine Th2- and Th1-biased responses respectively. The IAV NP mRNA vaccine induced higher IgG1 than IgG2c levels, while the IAV NP mRNA/DDO268 or DDO268B vaccine induced higher IgG2c levels, indicating a Th1-biased humoral response (**Fig. 6B** and **Fig. S2B**). These results align with our findings of enhanced IgG2c antibodies to an inactivated vaccine when DDO268 was added as an adjuvant [11, 12].

**Figure 6:**
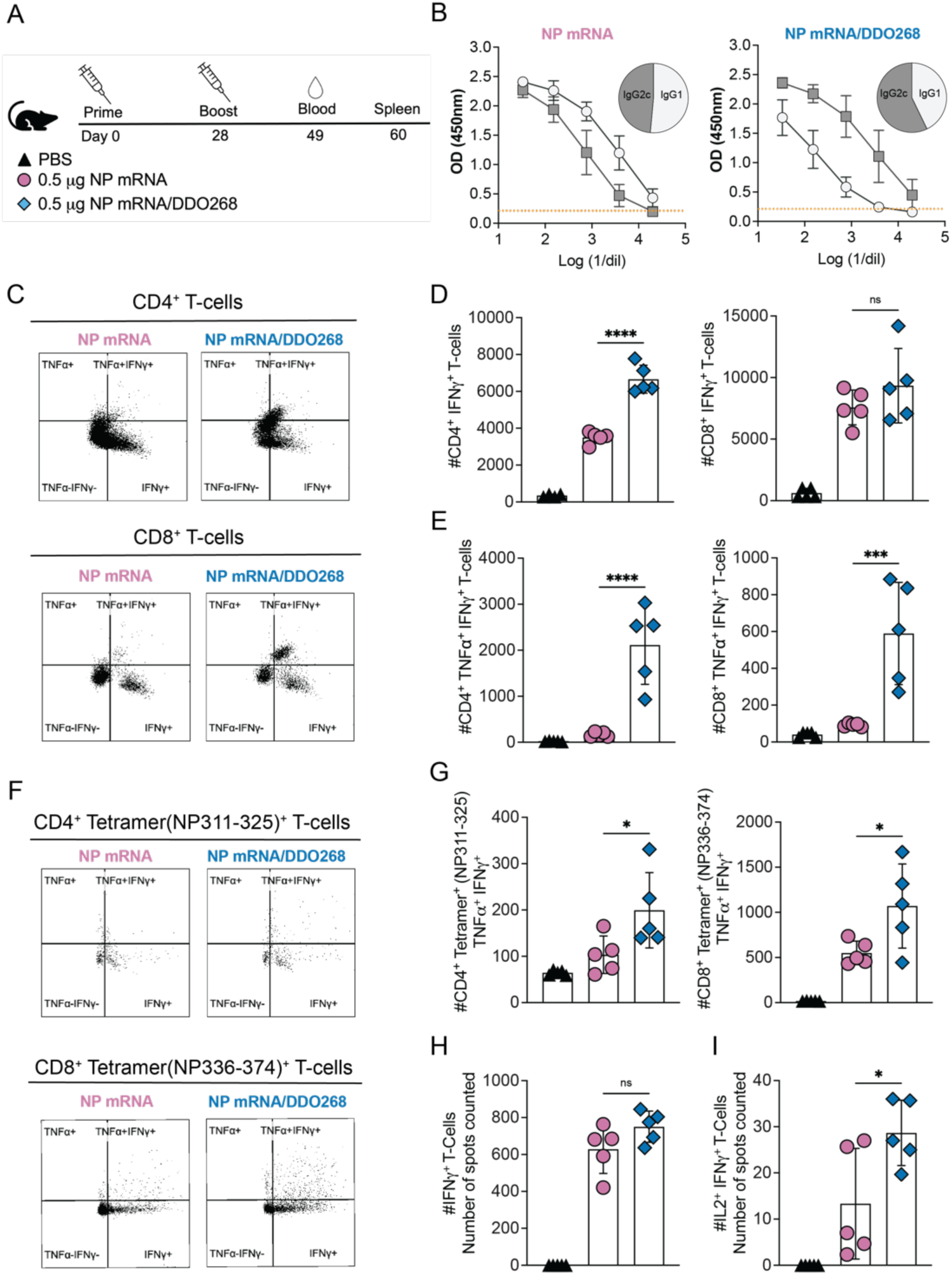
DDO268 promote strong type 1 adaptive humoral and cellular immune responses to the IAV NP mRNA vaccine. (**A**) Timeline and groups for the study design. C57BL6 mice were immunized twice 28 days apart with 0.5 µg NP mRNA or 0.5 µg of NP mRNA/DDO268. **(B)** Blood was collected 3 weeks after booster and specific NP antibodies IgG1 and IgG2c subtypes were evaluated by ELISA. Sera were serially diluted, and the orange line correspond to the cut off (normal mouse serum OD + 2DS). The pie graphs represent the ratio of IgG1 and IgG2c in mouse serum at a dilution of 1:32. **(C-I)** Antigen-experienced cells in the spleen were examined on day 32 after the booster immunization. **(C)** Representative flow cytometry plots for CD4^+^ and CD8^+^ TNFα^+^ IFNγ^+^ from the spleens of vaccinated mice. CD4^+^ TNFα^+^ IFNγ^+^ were identified by gating on live, singlets, CD3^+^ CD4^+^ CD8^−^ CD11a^+^ cells. CD8^+^ TNFα^+^ IFNγ^+^ were identified by gating on live, singlets, CD3^+^, CD8^+^ CD4^−^ CD11a^+^ cells. **(D)** Number of CD4^+^ IFNγ^+^ and CD8^+^ IFNγ^+^ T-cells in the spleens of individual mice in each vaccination group after Ionomycin/PMA restimulation. **(E)** Number of CD4^+^ TNFα^+^ IFNγ^+^ and CD8^+^ TNFα^+^ IFNγ^+^ T-cells in the spleens of individual mice in each vaccination group after Ionomycin/PMA restimulation. **(F)** Representative flow cytometry plots for CD8^+^ Tetramer (NP336-374)^+^ and CD4^+^ Tetramer (NP311-325)^+^ TNFα^+^ IFNγ^+^ T-cells. Specific CD4^+^ TNFα^+^ IFNγ^+^ were identified by gating on live, singlets, CD3^+^, CD4^+^ CD8^−^, Tetramer (NP311-325)^+^. CD8^+^ TNFα^+^ IFNγ^+^ were identified by gating on live, singlets, CD3^+^, CD8^+^ CD4^−^, Tetramer (NP336-374)^+^. **(G)** Number of tetramer specific CD4^+^ Tetramer (NP311-325)^+^ TNFα^+^ IFNγ^+^ and CD8^+^ Tetramer (NP336-374)^+^ TNFα^+^ IFNγ^+^ in the spleens of individual mice in each vaccination group after specific IAV NP restimulation. Number of cells shown was normalized to 500,000 live cells. **(H)** Number of spots counted as IFNγ^+^ after specific IAV NP restimulation (from 200,000 cells). **(I)** Number of spots counted as double-positives for IL2 and IFNγ after specific IAV NP restimulation (from 200,000 cells). Data corresponds to individual mice with mean ± SEM (n = 5/group). *=p <0.05, ***= p <0.005, ****= p <0.001 by one-way ANOVA with Bonferroni’s *multiple comparison* test.

To assess T cell responses, mice were euthanized 32 days post-booster immunization and their splenocytes were *in vitro* restimulated with Ionomycin/PMA. Both mRNA vaccines induced CD4^+^IFNγ^+^ and CD8^+^IFNγ^+^ T-cells in the spleen (**Fig. 6C** and **D**). However, the DDO268 adjuvanted NP mRNA vaccine induced significantly increased numbers of double-positive (TNFα^+^IFNγ^+^) CD4^+^ and CD8^+^T-cells (**Fig. 6C** and **E**) suggesting that DDO268 facilitates T cell activation to perform effector functions. Given that IAV NP contains T cells epitopes [25, 26], we also evaluated the induction of IAV-specific CD4^+^ T-cells and CD8^+^ T-cells. Both IAV NP mRNA and IAV NP mRNA/DDO268 or DDO268B formulations induced NP-specific CD8^+^ and CD4^+^ T-cells in the spleen. After *in vitro* IAV NP restimulation, the DDO268-adjuvanted mRNA vaccine induced significantly higher populations of IAV-specific CD8^+^ Tetramer (NP336-374)^+^ double positive for TNFα^+^IFNγ^+^ and CD4^+^ tetramer (NP311-325)^+^ double-positive for TNFα^+^IFNγ^+^ (**Fig. 6F** and **G**). Additionally, a 2-colors (IFNγ and IL2) ELISPOT assay showed higher double-positive spots for IFNγ and IL2 (**Fig. 6I** and **Fig. S2F**) in the DDO268 (or DDO268B)-containing formulation after *in vitro* restimulation. Overall, DDO268-adjuvanted formulations induced CD8^+^ and CD4^+^ T-cells characterized by both IFNγ and TNFα or IL2 production.

### IAV NP mRNA/DDO268 vaccine protects mice against IAV challenge

We next evaluated the protective efficacy or our mRNA vaccines against challenge with a lethal dose of IAV PR8 (H1N1). Thirty-nine days after the booster immunization, mice were challenged intranasally with 40 TCID_50_ of IAV PR8 and monitored for weight loss for 20 days **(Fig. 7A)**. In two independent experiemnts mice vaccinated with mRNA vaccines survived and recovered from the challenge. Notably, mice inoculated with the IAV NP mRNA/DDO268 vaccine had a higher survival probability after a lethal dose of virus (**Fig. 7B**). Although all mice experienced bodyweight loss (**Fig. 7C**), the DDO268-containing formulation induced a robust specific type 1 immune response, effectively protecting against IAV infection. Viral RNA transcripts were assessed 7 days post challenge (**Fig. 7D**) with *IAV NP* transcripts detected in both groups but reduced in mice vaccinated with IAV NP mRNA/DDO268. At the conclusion of the experiments, mice were sacrificed, and the presence of IAV NP-specific memory effectors CD8^+^ and CD103^+^ CD8^+^ T-cells in the lungs was evaluated. As shown in **Fig. 7E and F**, both vaccines generated a population of effector memory CD8^+^ tetramer^+^ (**Fig. 7E**) and CD8^+^ tetramer^+^ CD103^+^ (**Fig. 7F**). However, the percentage of these cells was higher in mice vaccinated with IAV NP mRNA/DDO268, indicating stronger protective immune response elicited by the DDO adjuvanted vaccine.

**Figure 7:**
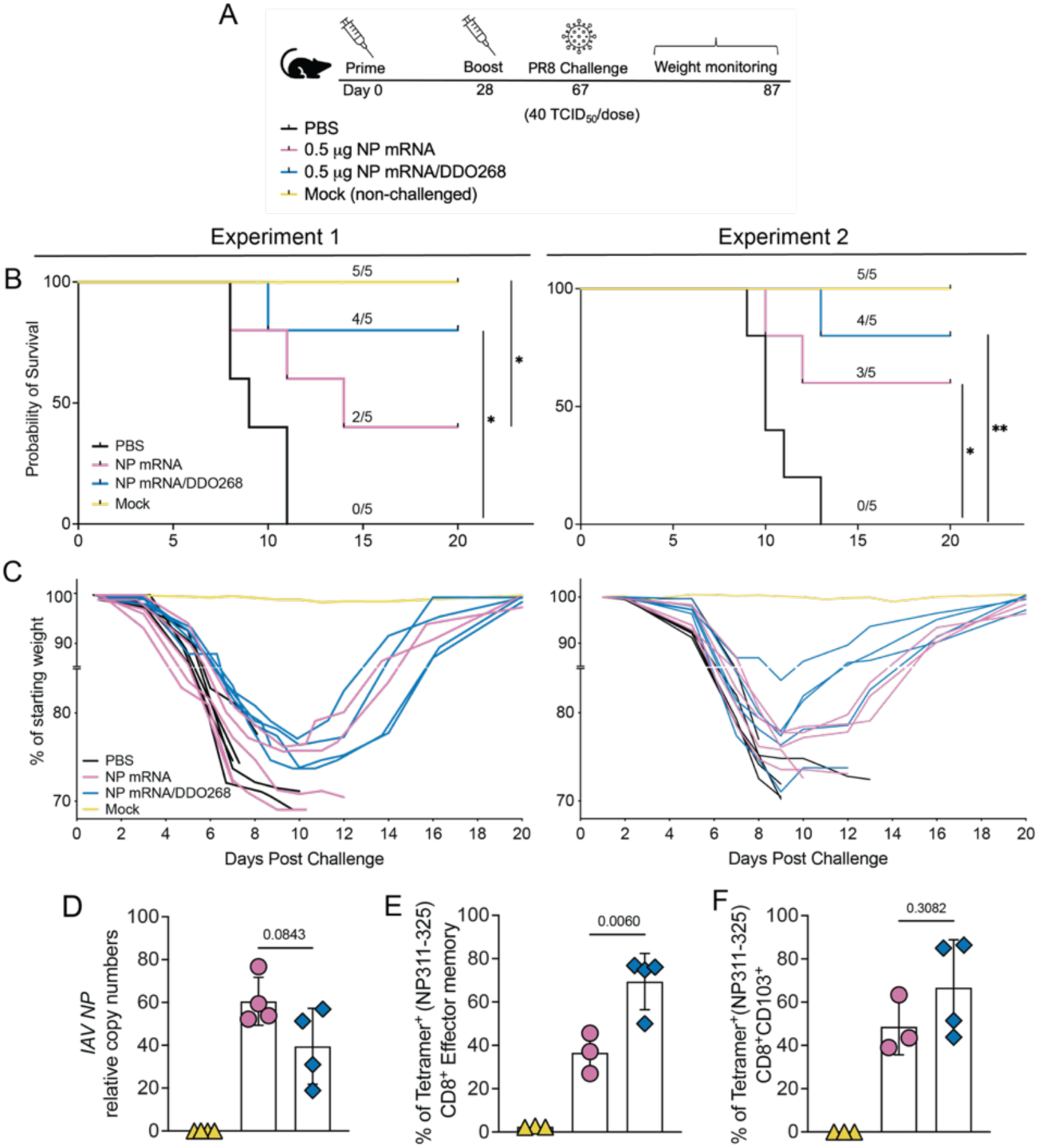
DDO268 enhances protective immunity against Influenza A/PR8 following IAV NP mRNA vaccination. **(A)** Timeline and groups for the study design. Vaccinated C57BL6 mice were challenged with 40 TCID_50_/dose of mouse passaged PR8 (H1N1) intranasally on day 39 after boost immunization. **(B)** Probability of survival in two independent experiments, n=5 mice each **(C)** weight relative to initial body weight over time in two independent experiments, n=5 mice each **(D-F)** Data corresponds to Experiment 2 n=3-5 (surviving animals). **(D)** Transcript levels of *IAV NP* relative to the housekeeping genes GAPDH and *β-actin* in the lungs of vaccinated mice at 7 days after challenge. **(E)** Percentage of effectors CD8^+^ in the lungs that are tetramer (NP336-374)^+^ at day 24 post challenge determined by staining and flow cytometry analysis. Effector Tetramer+ cells were identified after gating on single cells, live, CD3^+^, CD8^+^, CD44^+^, CD62L^-^, CCR7^-^, Tetramer^+^. **(F)** Percentage of CD8^+^ CD103^+^ T-cells in the lungs that are tetramer (NP336-374)^+^ at day 24 post challenge determined by staining and flow cytometry analysis (identified after gating on single cells, live, CD3^+^, CD8^+^, CD44^+^, CD103^+^, Tetramer(NP336-374)^+^. *= p <0.05 *<u>*</u> = p <0.01, as determined by log-rank Mantel-Cox test for survival and by one-way ANOVA with Bonferroni’s multiple comparison test.

## DISCUSSION

A significant challenge in developing vaccines that protect against intracellular pathogens is the lack of appropriate and safe adjuvants to drive robust type 1 cellular immunity, including both Th1 CD4^+^ T-cells and cytotoxic CD8^+^ T-cells. The ideal type 1 adjuvant would safely stimulate a broad range of cell types. Our lab identified the non-standard viral genome-derived RNA oligonucleotide, DDO268, that induces type I IFNs and proinflammatory cytokines by stimulating the cellular sensors RIG-I [20] and TLR3 [12]. DDO268 is a synthetic and replication-incompetent RNA derived from the 546-nucleotide-long Sendai virus (SeV) non-standard viral genome (nsVG), which is a primary immunostimulatory molecule during SeV infections [27, 28].

Our current results together with earlier research [12] provide robust evidence that the *in vivo* administration of DDO is safe and elicits a localized immune response without systemic effects (**Fig 1**), even at significantly higher doses than those used in previous studies [11, 12]. These findings enhance the safety profile of DDO and its potential as a valuable adjuvant in immunotherapeutic applications.

Our previous data [12] and initial experiments combining the BNT162b2 mRNA vaccine with DDO268 revealed that DDO268 effectively increases the migration and activation of conventional cDC1 in the draining lymph nodes, a critical step for initiating a robust type 1 immune response [29]. This increase in cDC1 numbers is not observed when BNT162b2 mRNA vaccine is combined with a non-immunostimulatory RNA. Additionally, the combination of BNT162b2 and DDO268 resulted in higher titers of IgG2c antibodies and a greater population of SARS-CoV-2 specific CD8^+^ IFNγ^+^ T-cells supporting DDO268 potential to enhance mRNA vaccines efficacy by promoting both humoral and cellular immune responses.

We have previously reported that when DDO is simply mixed with a purified protein antigen or an inactivated virus vaccine, it triggers the endosomal and cell surface receptor TLR3 [12]. TLR3 which is expressed in antigen-presenting cells, recognizes RNA, and signals for type I IFN production. However, when DDO is delivered intracellularly, it activates RIG-I-like receptors [20, 30]. One significant advantage of RIG-I stimulation is its universal expression in all animals and most nucleated cells, increasing the chances that the adjuvant activity of DDO will be conserved across larger animals and humans. Effectively targeting RIG-I requires intracellular delivery of DDO in lipid nanoparticles so that the RNA is deposited intracellularly. mRNA vaccines serve as an efficient carrier for this purpose.

mRNA vaccines can target a wide range of antigens, offering potential against diverse threats, including extracellular and intracellular pathogens and cancerous cells. Despite this, most mRNA vaccine efforts focus on antibody-mediated protection due to the crucial role of antibodies in neutralizing extracellular pathogens. However, antibodies are generally insufficient against intracellular pathogens or cancer. Current human influenza vaccines target the hemagglutinin protein (HA), which is not conserved across all strains. To develop a universal influenza vaccine, strategies have been explored to improve cross-reactive immunity against HA. One approach targets the more conserved stalk region of HA and has progressed to clinical trials. While these strategies induce antibodies that cross-react with multiple subtypes and protect against mortality, significant morbidity still occurs after infection (reviewed in [31, 32]). T cells recognize epitopes in the nucleoprotein (NP) of influenza viruses, which remains largely conserved despite antigenic shift, unlike the hemagglutinin and neuraminidase proteins that are the targets of neutralizing antibodies [33, 34].

In this study, we targeted the highly conserved NP in an influenza A mRNA vaccine and complemented it with DDO268 to elicit specific humoral and cellular responses capable of eliminating infected cells. We used unmodified dNTPs, cellulose-purified NP mRNA, demonstrating high stability and translation efficiency both *in vitro* **(Fig. 3C)** and *in vivo* **(Fig. 4C** and **D)**. To minimize type I interferon (IFN) induction from the mRNA in the vaccine, we incorporated a cellulose purification step, ensuring that DDO was the primary source of type I IFN in the formulated vaccine. Our results showed that cellulose purification significantly reduced type I interferon levels induced by IAV NP mRNA in cultured cells **(Fig. 3D)** and at the inoculation site **(Fig. 5C)**. These findings confirm that our unmodified dNTPs, cellulose-purified mRNA is effective and safe for use. While methyl-pseudouridine is commonly used during IVT to enhance mRNA stability and reduce immune responses [35] it can potentially lead to translational errors or “slippage.” causing aberrant proteins and immune recognition [36].

Lipid nanoparticles (LNPs) act as adjuvants for mRNA vaccines, enhancing the immune response by inducing the production of cytokines and chemokines. Previous studies have demonstrated that LNPs stimulate the production of *Il6, Il1b* and *Ccl2* [7] which recruit immune cells to the inoculation site. However, production of type I IFN is primarily driven by the mRNA [7]. Our results corroborate these findings, as we observed significant induction of *Il6, Il1b,* and *Ccl2* but no induction of *Ifnb1, Cxcl10,* or *Il13* when empty LNPs were used. Additionally, we observed minimal local induction of *Ifnb1* during immunization with IAV NP mRNA and IAV NP mRNA/X region **(Fig. 5C)** compared to the IAV NP mRNA/DDO268 formulation, demonstrating that most IFN induction is attributed to DDO268. These results highlight the advantages of our DDO-containing formulation which includes a controlled, local, and transient type I IFN induction, as shown in our **Fig. 5C** and **Fig. 5I-L** and supported by our previous studies [12].

Our current data, agrees with our previous findings [12] showing that the type 1 immune response induced by DDO268 in mice requires localized type I IFN production and the migration of conventional dendritic cells type 1 (cDC1) to the draining lymph nodes. cDC1 cross-present antigens playing a critical role in generating CD8^+^ T-cell responses to subunit and killed vaccines [37]. Here we demonstrate that the IAV NP mRNA/DDO268 vaccine enhances cDC1 activation and migration, subsequently priming CD8^+^ T-cells. DDO268-adjuvanted formulation also increases IgG2c levels in vaccinated mice **(Fig. 6B)**. This finding aligns with our previous reports [11, 12] where DDO268 enhanced IgG2c production in protein and purified virus vaccines. The higher IgG2c levels compared to the unadjuvanted vaccine suggests a skewing of the immune response towards a type 1 profile. Consistently, the DDO268 adjuvanted vaccine induced a higher population of fully activated NP-specific T-cells, as evidenced by increased IFNγ^+^TNFγα^+^ or IFNγ^+^IL2^+^ production **(Fig 6)**. This enhanced activation profile highlights the effectiveness of DDO268 in boosting the cellular immune response, crucial for long-term protection against viral infections.

The enhanced immune response generated by our DDO268-adjuvanted mRNA vaccine is evidenced by increased survival of mice following a lethal IAV PR8 infection (**Fig. 7**). This robust protection was observed even at a lower vaccine dose (0.5 µg) compared to the standard 5 µg dose. Our findings suggest that the DDO268 adjuvant not only enhances the immune response towards type 1 immunity but also allows to reduce the antigen dose required for protection, contributing to the vaccine effectiveness against IAV.

Overall, this study highlights the potent adjuvant effect of DDO268 in enhancing type 1 immune responses in SARS-CoV-2 and IAV mRNA vaccines. The ability of DDO268 to promote robust T-cell and humoral immunity, along with its efficacy in viral protection, emphasizes its potential as a valuable component in mRNA vaccine formulations. Future research will explore the application of DDO268 in other vaccine platforms for infectious diseases and cancer.

## MATERIAL AND METHODS

### Ethics statement

All described studies adhered to the Guide for the Care and Use of Laboratory Animal of the National Institute of Health. Institutional Animal Care and Use Committee, Washington University in St. Louis approved protocol 23-0083.

### Mice

C57BL/6 mice were obtained from The Jackson Laboratory. *Ifnar*1-/- mice were provided by Dr. Thomas Moran (Icahn School of Medicine at Mount Sinai) [38] and were used with sex and age matched C57BL/6 mice (Jackson Laboratory bred in house). Both male and female mice were included in the experiments.

### Viruses

Influenza A/Puerto Rico/8/1934 H1N1 (IAV PR8 H1N1) was used as challenge strains. IAV were grown in 10 day-old embryonated chicken eggs (Charles River Laboratory) at 30,000 TCID_50_ at 37 °C. Allantoic fluid from infected eggs was collected 40 hours later.

### mRNA production

The mRNA construct encodes for the nucleotide sequence of the NP of IAV A/Puerto Rico/8/1934 H1N1 (GenBank: ACV49549.1). The codon optimized sequence, synthesized by Twist Bioscience was cloned into the mRNA production plasmid (pJB201.1). This plasmid contains a T7 type II promoter, a 5’ UTR, a Kozak sequence with an AG mutation allowing for co-translational 5’ capping using CleanCap. The 5’ untranslated region (UTR) is from beta globin while the 3’ UTR is from alpha globin followed by a 142 base pair poly(A) tail. mRNA was produced using T7 RNA polymerase after completely linearization and capped co-transcriptionally with a trinucleotide cap-1 analog (TriLink) via the Hi Scribe T7 Clean Cap NEB kit. No chemically modified nucleosides were used. After IVT, mRNA size and degradation were assessed using Agilent Bioanalyzer 2100. dsRNA contaminants were removed following Baiersdorfer [9] protocol.

### mRNA construct testing

The IVT mRNA was tested in HEK-293t and A549 cells using jetMESSENGER™ (Polyplus) as transfection reagent. IAV NP production from the IVT mRNA was measured 24 and 48 hours after transfection by intracellular staining using NP IAV antibody NBP3-12741AF647 (Novus Biological) followed by flow cytometry.

### Stimulatory RNA production

DDO268, DDO268B and the control X region expressing plasmid (with a T7 promoter) were linearized and *in vitro* transcribed with the Hi Scribe T7 (NEB). RNA products were DNase-treated and precipitated with LiCl. The OD260/OD280 ratios were between 2.00 and 2.25, and the OD260/OD230 ratios were between 2.20 and 2.60. RNA purity and integrity were confirmed using an Agilent Bioanalyzer 2100 with endotoxin level below 0.1EU/ml/300ug.

### Vaccine formulation

Formulations of NP mRNA, NP mRNA/X region, NP mRNA/DDO268, and NP mRNA/DDO268B, were encapsulated in a lipid nanoparticle, GenVoy Ionizable Lipid Mix (ILM), and using NanoAssemblr Ignite (Precision Nanosystems) following manufacture protocol. Encapsulation efficiency and concentration of the RNA were tested by Ribogreen Assay (ThermoFisher) following manufacturer protocol. The final particle size was tested using DLS. mRNA and adjuvant RNA co-packaging was performed at 1:1 molar ratio, ensuring that each formulation contained the same total amount of RNA (about 0.5 ug total RNA).

### Mice immunization

Mice were anesthetized with isoflurane and injected subcutaneously (s.c.) into a rear footpat. For toxicity studies with 50 µg of DDO268 or PBS at a final volume of 30 µl per dose. For immune response studies mice were inoculated with BNT162b2 (0.125 µg); BNT162b2 (0.125 µg) + DDO268 (5 µg); BNT162b2 (0.125 µg) + RNA (5 µg); 1ug IAV NP mRNA; 1 µg IAV NP mRNA/DDO268; 0.5 µg IAV NP mRNA; 0.5 µg IAV NP mRNA/X region; 0.5 µg IAV NP mRNA/DDO268; 0.5 µg IAV NP mRNA/DDO268B; Empty LNPs or PBS at a final volume of 30 µl per dose. The BNT162b2 SARS-CoV-2 vaccine was leftover vaccine that was collected and stored at -80°C again prior to use in this study. For innate immune experiments mice were vaccinated once and for the adaptive immune response experiments mice were primed and boosted 28 days later with the same vaccine formulations.

### Mice challenge

Mice were challenged intranasally with 40 TCID_50_ of IAV-PR8 39 days after boost. All mice were weighed daily post-challenge and survival probability was recorded based on an increase or decrease of mortality.

### Complete Blood Count

Peripheral blood samples were collected from mice via cardiac puncture and blood was collected into EDTA-coated tubes to prevent clotting. Samples were processed immediately and analyzed using an automatic hematology analyzer. The results were generated automatically by the analyzer software and recorded.

### Blood chemistry analysis

Peripheral blood samples were collected from mice via cardiac puncture and collected into serum-separating tubes. Samples were processed immediately following standard protocols for each parameter.

### RT-qPCR

Transfected A549 cells, mice footpads, spleen liver and lungs were harvested and total RNA was extracted using TRIzol (Ambion Inc.) following manufacturer’s guidelines followed by DNAse I (Thermo scientific) treatment. cDNA synthesis was performed with the High-Capacity cDNA Reverse Transcription kit (Applied biosystems). For quantitative analysis by RT-PCR (qPCR), 10 ng/µL of cDNA was amplified using SYBR Green Mastermix (Thermofisher) in a BioRad C1000 Touch thermal cycler (BioRad). Transcripts were measured using the delta-delta Ct method. Primers used are listed in **Table 1**.

**Table 1:**
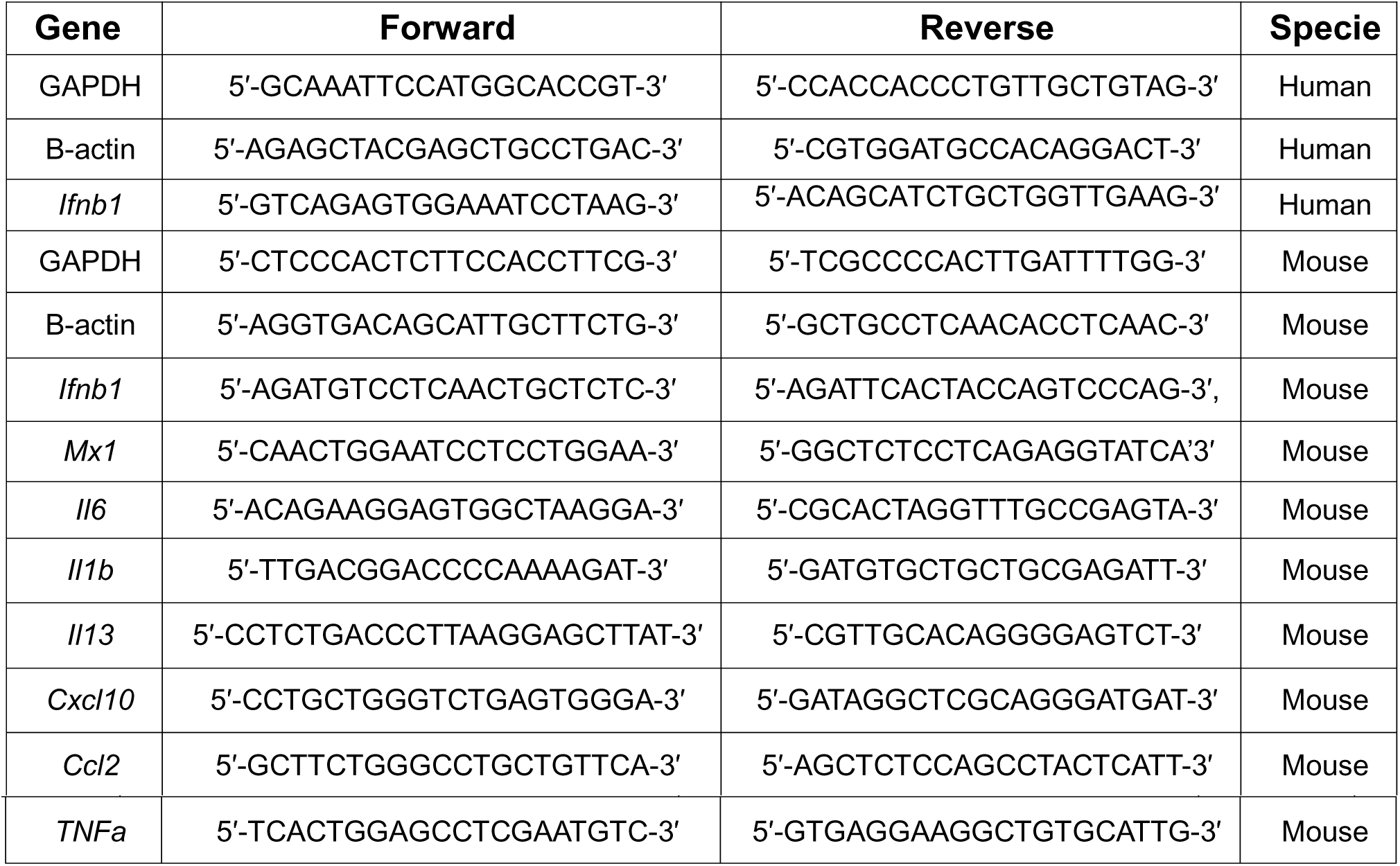
Primer sequences used for quantitative analysis by RT-PCR (qPCR).

### Single-cell suspension and flow cytometry staining

<u>Footpads</u> were processed following Fangzhou et al [39] with modifications. Digested samples were vortexed and filtered through a 70-µm mesh to obtain single-cell suspensions. Immune cells were defined using a panel of surface markers. <u>Lymph nodes</u> were digested using DNase (1 µg/ml) and Liberase (5 µg/ml) in RPMI 1640 Medium (Life Technologies) for 20 min at 37 °C, then vortexed and filtered through a 70-µm mesh to obtain single-cell suspensions. cDC1 cells were identified using a surface marker panel, and IAV expression was tested by intracellular staining. <u>Spleens</u> were collected in RPMI 1640 Medium (Life Technologies) and filtered through a 70-µm mesh to obtain single-cell suspensions. Cells were treated with red blood cells lysis buffer (Sigma). For each sample, 1×10^6^ cells were restimulated using PMA (0.1 mg·mL) and Ionomycin (1mg/ml) or purified IAV NP protein (Sino biological) for 4 h at 37 °C followed by Brefeldin treatment. Cells were resuspended in PBS with 1% FBS (Sigma) and blocked with anti-mouse FcγRIII/II (CD16/32; BD Biosciences) for 20 min on ice. T cell subpopulations were defined using a surface marker panel, with activation determined by intracellular staining for IFNγ and TNFα. For intracellular staining, cells were fixed/permeabilized with the FoxP3/Transcription Factor Staining Buffer Set (eBioscience) and incubated with anti-IAV NP antibody conjugated with AlexaFluor-647 (Invitrogen). <u>Lungs</u> were inflated with 0.7 ml of digestion mix containing collagenase A (Sigma), dispase (Thermofisher), Liberase TL (Sigma) and DNAse I (Sigma) and incubated at 37°C for 30 min with agitation. Digested samples were filtered through a 70-µm filter mesh to obtain single-cell suspensions. The obtained cells were washed with PBS containing 5% FBS, treated with red blood cells lysis buffer (Sigma), and total viable cells were quantified with trypan blue staining using an automated cell counter (TC-20 Automated Cell Counter; BioRad).

### Spectral flow cytometry

Flow cytometry experiments were performed using a Cytek Aurora spectral flow cytometer (Cytek Biosciences), with at least 1×10^6^ events acquired. Data were analyzed using FlowJo V12 software (Tree Star Inc.). Single-cell suspensions were stained with fluorochrome-labeled antibodies. Fixable Viability Dye eFluor506, monoclonal antibody specific for mouse IFNγ (XMG1.2), Ly6c (HK1.4), CD3 (17A2), CD19 (eBio1D3), B220 (RA3-6B2), NK1.1 (PK136), CD11b (M1/70), and TNFα (MP6-XT22) were obtained from eBioscience. Monoclonal antibodies specific for mouse CD3 (17A2), CD4 (GK1.5), CD11a (H11578), XCR1 (ZET), PDCA1 (129 cl), CD11c (cN418), SIRPα (P84), CD64 (x54/7.1), CD44 (NIM-R8), CD62L (cMEL-14), CCR7 (4B12), CD103 (QA17A24) and MHC-II (M5/114.15.2) were obtained from Biolegend. Monoclonal antibodies for mouse CD86 (GL1) and CD8 (53–6.7) were obtained from BD BioSciences. IAV-specific tetramers: H-2D^b^ tetramers bearing NP366-374 (ASNENMETM) and I-A^b^ tetramers bearing NP311-325 (QVYSLIRPNENPAHK) were obtained from NIH Tetramer Core Facility at Emory University. APC-labeled SARS-CoV-2 S-specific tetramer (MHC class I tetramer, residues 539–546, VNFNFNGL, H-2Kb).

### Quantification of influenza NP-specific serum antibodies

Sera from 21 post-boost immunized mice were analyzed for anti-SARS-CoV-2 Spike or anti-IAV NP IgG1 and IgG2c antibodies. ELISA plates (Nunc, MaxiSorp) were coated with 2 µg of purified SARS-CoV-2 Spike (Kindly provided by Dr. Ellebedy, Washington University in St. Louis) or IAV-NP and treated with pre-diluted sera in triplicate, followed by HRP-conjugated anti-mouse IgG1 or IgG2c (Sourthen Biotech) and TMB substrate (Sera Care).

### IFN-**γ**, IL2 ELISPOT Assays

Double-color fluorescent ImmunoSpot kits from CTL were used for the detection of in vivo-primed IFN-γ-IL2 producing Th1-type memory T-cells. In brief, IAV NP or Ionomycin/PMA were plated at the specified concentrations into capture antibody-precoated ELISPOT assay plates. 200,000 viable cells/well in 100μL CTL-Medium and cultured with the NP or mitogen for 24h at 37 °C and 9% CO_2_. After addition of detection antibody, and enzymatic visualization of plate-bound cytokine, the plates were scanning, counting of spot forming units and analyzed using an S6 Ultra M2 Fluorospot reader, by CTL. Spot numbers were automatically calculated by the ImmunoSpot Software using the Autogate function of the ImmunoSpot Software.

### Virus titration

To obtain lung viral titers, lung lobes were homogenized in 0.1% Gelatin clarified by centrifugation, and analyzed by infectivity assays in MDCK cells. Virus titers were determined using end-point dilution tissue culture infectious dose (TCID50) infectivity assay (Reed and Muench).

### Statistical analysis

Statistical significance was inferred using GraphPad Prism software version 10.0 (GraphPad Software, San Diego, CA). Unpaired t-test, one-way and two-way ANOVA with Bonferroni’s multiple comparison test. Log-rank Mantel-Cox test for survival. For animal experiments, group size consisted of 3-5 mice per group.

## Authors contributions

Conceived experiments: V.G., and C.B.L.; performed experiments and collected data: V.G, H.S, Wrote the original draft: V.G. and C.B.L.; Supervised research activities: C.B.L. Design and plasmid validation A.K.P and J.D.B

## Supporting information

All supplementary figures

## Acknowledgements

The authors wish to thank: Dr. Michael Diamond and Pritesh Desai (Washington University in St. Louis) for providing SARS-CoV-2 tetramer, Dr. Ali Ellebedy (Washington University in St. Louis) for providing SARS-CoV-2 Spike protein and ELISA protocol. This work was supported by the US National Institutes of Health National Institute of Allergy and Infectious Diseases (NIH R01AI134862 to C.B.L. R01AI1506701A1 to J.D.B) and 3U01CA260541-02S1 to J.D.B.

## Data availability

All data are available upon request.

