## Supplementary material for "DDO-adjuvanted influenza A virus nucleoprotein mRNA vaccine induces robust humoral and cellular type 1 immune responses and protects mice from challenge": All supplementary figures

**FIGURE S1**

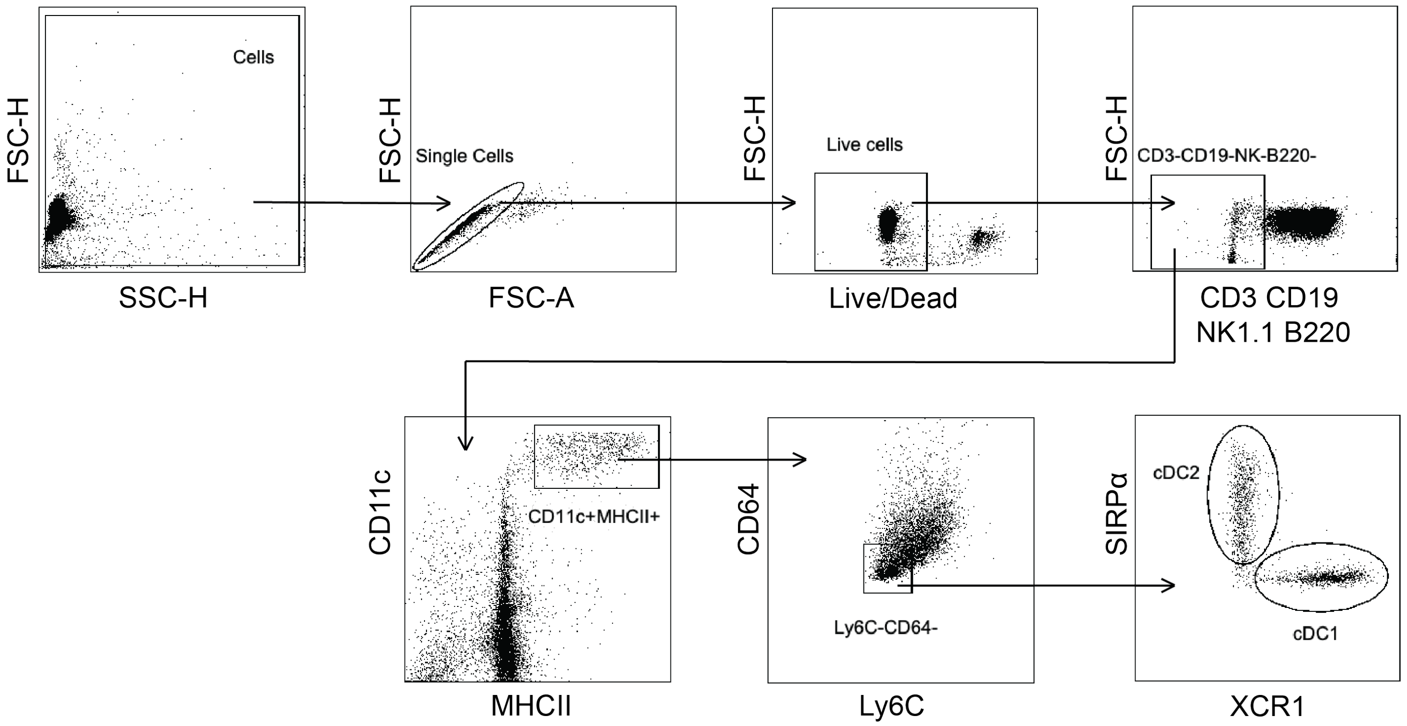


**Figure S1. Gating strategy for spectral flow cytometry analysis of conventional dendritic cells type 1 (cDC1).** Representative dot plots indicating the following cell subsets defined from live cells: Non-Lymphoid cells ($CD3^{-}NK1.1^{-}CD19^{-}B220^{-}$), Dendritic cells ($CD11c^{hi}MHCII^{hi}$), Conventional Dendritic cells ($Ly6G^{-}CD64^{-}$), cDC1 ($XCR1^{+}Sirpa^{-}$), cDC2 ($Sirp\alpha^{+}XCR1^{-}$).

FIGURE S2

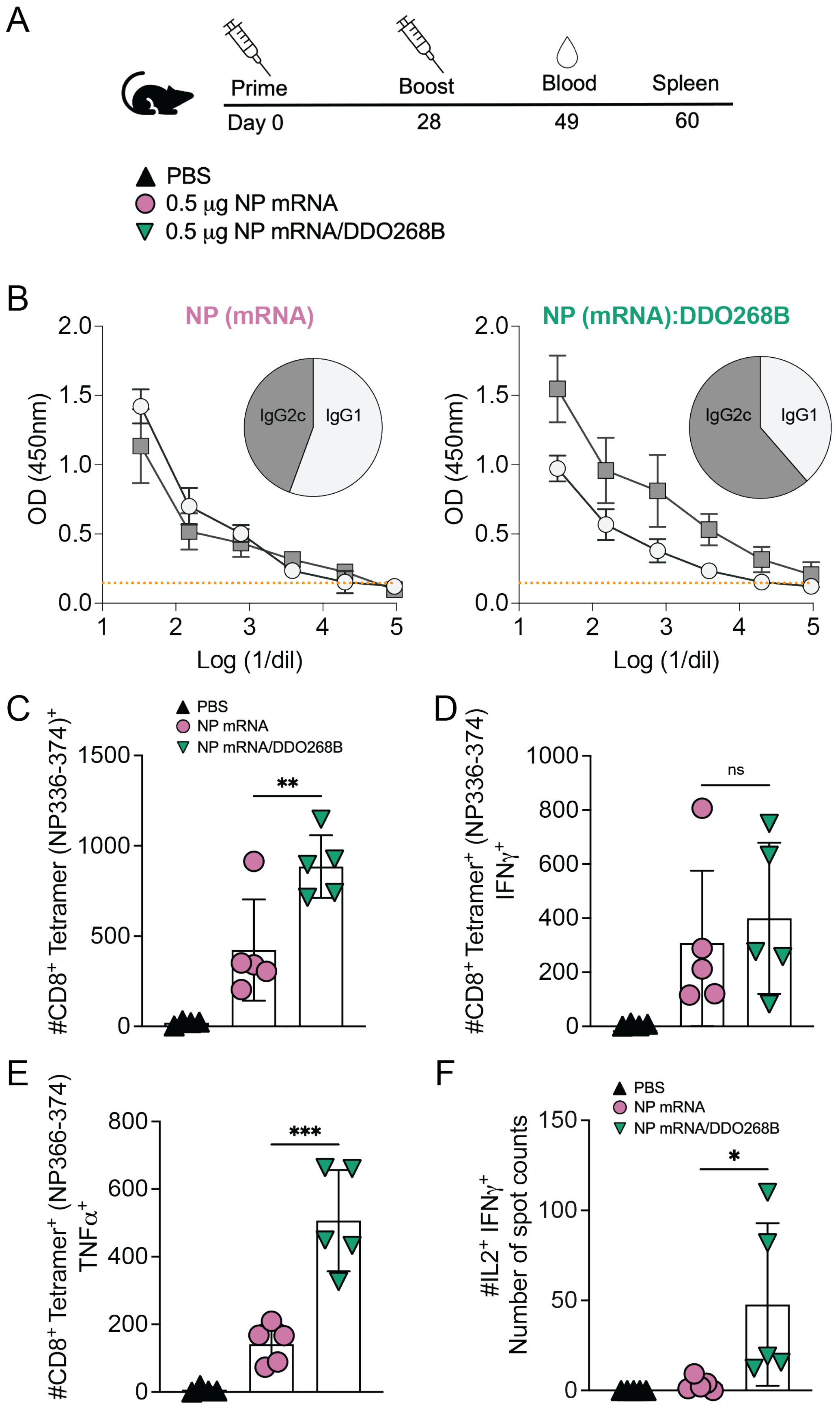
A
Prime
Boost
Blood
Spleen
Day 0
28
49
60
PBS
0.5 µg NP mRNA
0.5 µg NP mRNA/DDO268B
B
NP (mRNA)
NP (mRNA):DDO268B
IgG2c
IgG1
OD (450nm)
Log (1/dil)
C
#CD8+ Tetramer (NP336-374)+
D
#CD8+ Tetramer+ (NP336-374) IFNγ+
ns
E
#CD8+ Tetramer+ (NP366-374) TNFα+
F
#IL2+ IFNγ+ Number of spot counts
PBS
NP mRNA
NP mRNA/DDO268B

**Figure S2**: **DDO268B promote enhanced type 1 adaptive humoral and cellular immune responses to the IAV NP mRNA vaccine.** (**A**) Timeline and groups for the study design. C57BL6 mice were immunized twice 28 days apart with 0.5 µg NP mRNA or 0.5 µg of NP mRNA/DDO268B. **(B)** Blood was collected 3 weeks after booster and specific NP antibodies IgG1 and IgG2c subtypes were evaluated by ELISA. Sera were serially diluted, and the orange line correspond to the cut off (normal mouse serum OD + 2DS). The pie graphs represent the ratio of IgG1 and IgG2c in mouse serum at a dilution of 1:32. Data corresponds to individual mice with mean ± SEM (n = 5/group). **(C-E).** Antigen-experienced cells in the spleen were examined on day 32 after the booster immunization. **(C)** Number of $CD8^+$ Tetramer $(NP336\text{-}374)^+$ T-cells in the spleens of individual mice in each vaccination group after specific *in vitro* IAV NP restimulation. **(D)** Number of $CD8^+$ Tetramer $(NP336\text{-}374)^+$ $IFN\gamma^+$ T-cells in the spleens of individual mice in each vaccination group after *in vitro* IAV NP restimulation. **(E)** Number of $CD8^+$ Tetramer $(NP336\text{-}374)^+$ $TNF\alpha^+$ T-cells in the spleens of individual mice in each vaccination group after *in vitro* IAV NP restimulation. $CD8^+$ $TNF\alpha^+$ or $CD8^+$ $IFN\gamma^+$ were identified by gating on live, singlets, $CD3^+$ $CD8^+$ Tetramer $(NP336\text{-}374)^+$. Number of cells shown was normalized to 500,000 live cells. **(F)** Number of spots counted as double-positives for IL2 and IFNγ after specific *in vitro* IAV NP restimulation (from 200,000 cells). Data corresponds to individual mice with mean ± SEM (n = 5/group). *=p <0.05, **= p <0.01, ***= p <0.005, by one-way ANOVA with Bonferroni's *multiple comparison* test.
