## Supplementary material for "DDO-adjuvanted influenza A virus nucleoprotein mRNA vaccine induces robust humoral and cellular type 1 immune responses and protects mice from challenge": Table S1

**Table 1:** CBC and chemistry profile of mice inoculated with DDO

| **CBC**  **PARAMETER** | **UNIT** | **ANIMAL ID** | | | | | | | | | |
| --- | --- | --- | --- | --- | --- | --- | --- | --- | --- | --- | --- |
|  |  | **DDO 24 h** | | | | **DDO 72 h** | | | | **PBS** | |
| **Leukogram** |  |  |  |  |  |  |  |  |  |  |  |
| WBC | K/uL | 5.56 | 4.54 | 6.50 | 4.50 | 7.92 | 7.48 | 8.38 | 13.24 | 7.38 | 3.46 |
| NE# | K/uL | 1.67 | 0.91 | 1.43 | 0.99 | 1.35 | 1.20 | 1.17 | 3.31 | 1.18 | 1.04 |
| LY# | K/uL | 3.50 | 3.41 | 4.62 | 3.38 | 6.34 | 6.06 | 6.96 | 9.27 | 6.05 | 2.25 |
| MO# | K/uL | 0.22 | 0.18 | 0.39 | 0.14 | 0.24 | 0.22 | 0.25 | 0.53 | 0.15 | 0.10 |
| EO# | K/uL | 0.17 | 0.05 | 0.07 | 0.00 | 0.00 | 0.00 | 0.00 | 0.13 | 0.00 | 0.07 |
| BA# | K/uL | 0.00 | 0.00 | 0.00 | 0.00 | 0.00 | 0.00 | 0.00 | 0.00 | 0.00 | 0.00 |
| NRBC# | K/uL | 0.00 | 0.00 | 0.00 | 0.00 | 0.00 | 0.00 | 0.00 | 0.00 | 0.00 | 0.00 |
| NE% | % | 30 | 20 | 22 | 22 | 17 | 16 | 14 | 25 | 16 | 30 |
| LY% | % | 63 | 75 | 71 | 75 | 80 | 81 | 83 | 70 | 82 | 65 |
| MO% | % | 4 | 4 | 6 | 3 | 3 | 3 | 3 | 4 | 2 | 3 |
| EO% | % | 3 | 1 | 1 | 0 | 0 | 0 | 0 | 1 | 0 | 2 |
| BA% | % | 0 | 0 | 0 | 0 | 0 | 0 | 0 | 0 | 0 | 0 |
| NRBC% | % | 0 | 0 | 0 | 0 | 0 | 0 | 0 | 0 | 0 | 0 |
| **Erythrogram** |  |  |  |  |  |  |  |  |  |  |  |
| RBC | M/uL | 11.45 | 10.42 | 10.64 | 10.43 | 10.63 | 10.77 | 11.47 | 13.46 | 10.69 | 10.25 |
| HB | g/dL | 14.6 | 13.0 | 13.9 | 13.8 | 14.1 | 14.2 | 14.4 | 17.8 | 14.7 | 13.8 |
| HCT | % | 51.8 | 48.6 | 50.3 | 49.1 | 50.5 | 50.0 | 54.5 | 63.0 | 50.6 | 48.1 |
| MCV | fL | 45.2 | 46.6 | 47.3 | 47.1 | 47.5 | 46.4 | 47.5 | 46.8 | 47.3 | 46.9 |
| MCH | Pg | 12.8 | 12.5 | 13.1 | 13.2 | 13.3 | 13.2 | 12.6 | 13.2 | 13.8 | 13.5 |
| MCHC | g/dL | 28.2 | 26.7 | 27.6 | 28.1 | 27.9 | 28.4 | 26.4 | 28.3 | 29.1 | 28.7 |
| RDW | % | 16.4 | 16.2 | 16.8 | 16.5 | 16.8 | 16.7 | 16.8 | 18.9 | 16.8 | 16.1 |
| **Thrombocytes** |  |  |  |  |  |  |  |  |  |  |  |
| PLT | K/uL | 354 | 690 | 759 | 497 | 477 | 560 | 968 | 920 | 512 | 492 |
| MPV | fL | 5.4 | 5.2 | 5.1 | 5.3 | 5.6 | 5.5 | 5.2 | 5.4 | 5.2 | 5.5 |
| PDW | % | 0 | 0 | 0 | 0 | 0 | 0 | 0 | 0 | 0 | 0 |

| **CHEMICAL**  **PARAMETER** | **UNIT** | **ANIMAL ID** | | |
| --- | --- | --- | --- | --- |
|  |  | **DDO 24 h** | **DDO 72 h** | **PBS** |

| AST | U/L | 466 | 143.0 | 108.0 | 77.0 | 187.0 | 100.0 | 69.0 | 46.0 | 213.0 | 590 |
| --- | --- | --- | --- | --- | --- | --- | --- | --- | --- | --- | --- |
| Blood Urea Nitrogen | mg/dL | 18.0 | 19.0 | 22.0 | 31.0 | 16.0 | 12.0 | 21.0 | 24.0 | 18.0 | 20.0 |
| Triglycerides | mg/dL | 85.0 | 95.0 | 177.0 | 89.0 | 69.0 | 70.0 | 97.0 | 97.0 | 93.0 | 107.0 |
| ALP | U/L | 130.0 | 140.0 | 143.0 | 120.0 | 146.0 | 134.0 | 117.0 | 150.0 | 167.0 | 124.0 |
| CK | U/L | 2022 | 534.0 | 425.0 | 359.0 | 791.0 | 332.0 | 214.0 | 116.0 | 956.0 | n/a |
| hsCRP | mg/L | 0.216 | 0.117 | 0.142 | 0.093 | 0.108 | 0.049 | 0.048 | 0.093 | 0.141 | 0.158 |
| LDH | U/L | 541.0 | 219.0 | 258.0 | 196.0 | 263.0 | 180.0 | 271.0 | 255.0 | 316.0 | 1520 |
| Total Bilurubin | mg/dL | 0.3 | 0.2 | 0.3 | 0.3 | 0.2 | 0.2 | 0.3 | 0.3 | 0.3 | n/a |
